# Polyploidy and plant-fungus symbiosis: evidence of cytotype-specific microbiomes in the halophyte *Salicornia* (Amaranthaceae)

**DOI:** 10.1101/2022.03.09.483717

**Authors:** Danilo Reis Gonçalves, Rodica Pena, Dirk C. Albach

## Abstract

Polyploidy is recognized as a mechanism of speciation in plants with cascading effects on biotic interactions. However, a limited number of studies have investigated the effects of polyploidy on the association of plants and microorganisms. Herein, we investigated whether two *Salicornia* cytotypes (*S. europaea* – 2x and *S. procumbens* – 4x) show different root-associated fungal communities. Additionally, we explored the existence of cytotype-specific root anatomical traits, which could influence fungal recruitment and establishment. *Salicornia* spp. were identified based on their ploidy level. The root-associated fungal microbiome of *Salicornia* was analyzed using high throughput amplicon sequencing (ITS1 of rDNA) in spring and summer. The following root anatomical traits were investigated: maximum root diameter, periderma thickness, parenchyma thickness, diameter of the vascular cylinder and maximum diameter of parenchyma cells. Our results showed that Shannon diversity and evenness indices were higher in samples of *Salicornia procumbens* (4x) compared to those of *S. europaea* (2x), and in summer the root-associated fungal community of *S. procumbens* (4x) was significantly different from that of *S. europaea* (2x). The orders *Xylariales, Malasseziales* and *Pleosporales* were the most frequent root colonizers in both cytotypes and most of the taxa associated with *Salicornia* were functionally classified as saprophytes or plant pathogens. Finally, we observed larger periderma and parenchyma layers in *S. procumbens* (4x) than *S. europaea* (2x) that may contribute to the observed differences in community composition between the two cytotypes. Our results suggest that differences in ploidy may modulate plant interaction with fungi by affecting species recruitment and microbiome structure. In addition, cytotype-specific root traits may also have the potential to affect differently community assembly in the two cytotypes.

## Introduction

Among the different factors controlling species interactions, polyploidy or whole-genome duplication is considered an important yet underexplored process (Segraves & Anneberg 2016). The genetic changes caused by polyploidization in plants may have effects on cell biology (Doyle & Coate 2019), plant chemistry (Corneillie et al. 2019; Veach et al. 2018), physiology (Baker et al. 2017) and anatomy (Chansler et al. 2016). These changes have been shown to produce cascading effects on biotic interactions, for example, on plant-pollinator (Husband and Schemske 2000; Kennedy et al. 2006) and plant-herbivore associations (Collins & Müller□Schärer 2012; Kao 2008). As an example, Edger et al. (2015) reported that the presence of glucosinolate defensive compounds in polyploid individuals of Brassicales might negatively impact herbivory. In the same way, an increase in the production of defense compounds in polyploids may negatively affect colonization by endophytic fungi (Segraves & Anneberg 2016; van Loon et al. 2006). However, associations of plants and microorganisms have been investigated in only a very limited number of studies, with controversial outcomes despite the prominent effect of microorganisms on the growth and survival of plants.

Investigations of potential effects of polyploidy on plant-microbe interactions are limited to bacteria (Cavé-Radet et al. 2020; Forrester & Ashman 2018, Forrester & Ashman 2019; Wipf & Coleman-Derr 2021) and arbuscular mycorrhizal fungi (AMF) (Annenberg & Seagraves 2019; Sudová et al. 2010; Sudová et al. 2014; Sudová et al. 2018; Tĕšitelová et al. 2013). For instance, microscopical analyses have shown that AMF colonization in tetraploid individuals of *Heuchera cylindrica* was higher compared to diploids (Anneberg & Segraves 2019) and more diverse AMF communities were observed in tetraploid individuals of *Gymnadenia conopsea* compared to diploids (Tĕšitelová et al. 2013). In contrast, two other studies showed no correlation of ploidy and AMF colonization in *Aster amellus* (Sudová et al. 2014) and *Centaurea stoebe* (Sudová et al. 2018). Importantly, none of these studies investigated the dynamics of colonization throughout different seasons. Thus, any general conclusions on the effects of polyploidy in shaping plant-fungus association in a broader context are currently not possible.

Despite the reported differences in microorganism colonization between diploid and polyploid plants, the underlying differences in anatomical, physiological or chemical processes directly affecting the plant-AMF interaction have not been explored in previous studies and are largely unknown. In the case of bacteria, this has started to be elucidated. Micallef et al. (2019) observed that in *Arabidopsis thaliana*, the different exudation patterns observed in different cytotypes directly influenced bacterial recruitment and community assembly, which could also be occurring with fungi. In addition, root anatomical traits likely play a role by affecting bacterial and fungal microbiome assembly. For example, Galindo-Castañeda et al. (2019) reported less mycorrhizal colonization in roots of individuals of *Zea mays* with a smaller living cortical area.

An interesting model organism to study the effects of polyploidy on species interactions is the halophyte *Salicornia*. In this genus, fungal symbionts, especially of the order *Pleosporales* (Ascomycota), are reported in the roots (Furtado et al. 2019a) and were shown to positively affect nutrient uptake and plant growth (Gonçalves et al. 2021). In northern Germany, two different but closely related *Salicornia* cytotypes occur, *Salicornia europaea* (2x) and *Salicornia procumbens* (4x). The two species have a similar morphology and react highly plastic to changes in the environment, which makes morphology-based species identification challenging and time-consuming. In addition to that, there are no studies investigating whether the two cytotypes differ in root anatomical traits, which could directly influence fungal recruitment and community establishment, as observed for endophytic bacteria (Forrester et al. 2020) and fungi (Galindo-Castañeda et al. 2019). Because of the morphological similarities of the cytotypes, species identification is mostly conducted based on their differences in ploidy (Kadereit et al. 2007). Although the broadest phylogenetic study of the genus *Salicornia* (Kadereit et al. 2007) suggested that both cytotypes are phylogenetically not sister, a recent study by Buhk (2020) using genotyping-by-sequencing across the western part of the German Wadden Sea, revealed a high genetic similarity and sharing of alleles between the cytotypes. Thus, the data by Buhk (2020) suggest an autopolyploid origin of *S. procumbens* by autopolyploidy from *S. europaea* or a genetically similar species.

Both cytotypes occur sympatrically but are differentiated on a microscale in the salt marsh zones. *Salicornia europaea* (2x) is mostly present in the lower salt marsh but also reported in the pioneer zone. On the other hand, *S. procumbens* (4x) seem to be more evenly distributed in the salt marsh zones and tidal flat, except in the upper salt marsh where both cytotypes do not occur. Interestingly, an ecologically differentiated distribution of two *Salicornia* species (*S. procumbens* and *S. stricta*) in the salt marsh zones was observed by Teege et al. (2011) with *S. procumbens* growing in the lowest parts of the salt marsh. The authors suggested a strong selection during seed germination and establishment as a reason for the existence of intraspecific ecotypes. Nevertheless, there is no information available on the role of belowground interactions regarding habitat differentiation between diploids and tetraploids, although it has been shown to influence habitat differentiation of plant species (Reynolds et al. 2003).

There are studies showing that in order to synthesize additional chromosomal sets, polyploids have a higher demand for nitrogen and phosphorus compared to diploids (Šmarda et al. 2013; Wildermuth 2010). This may have consequences for the association of plants with beneficial fungi. For example, Anneberg & Segraves (2019) observed a higher number of nutritional-exchange structures of AMF in roots of the tetraploid *Heuchera cylindrica* compared to diploids. In the case of *Salicornia*, in which AMF have been reported to be absent in the roots (Furtado et al. 2019a), polyploidy may have further consequences in plant’s interaction with another group of endophytic fungi, dark septate endophytes (DSE). Fungi classified as DSE are mainly found in the orders Pleosporales, Xylariales and Helotiales of the Ascomycota (Knapp et al. 2018), are commonly found in roots of salt marsh plants (Furtado et al. 2019a; Maciá-Vicente et al. 2016) and are known for their ability to improve plant growth (Mateu et al. 2020; Vergara et al. 2019). In this sense, it is possible that *S. procumbens* (4x) benefits from the association with a more diverse fungal community, especially of mutualistic DSE, to survive in the lowest parts of the salt marsh. In addition, the fitness benefits conferred by DSE in halophytes, such as salt stress tolerance (Mateu et al. 2020), may contribute to the adaptation of *S. procumbens* (4x) to more stressed areas (e.g., tidal flats), where frequent flooding associated with high salinity occur. Plant phenology is a major driver of plant ability to acquire soil resources (Nord & Lynch, 2009). Therefore, the fungal contribution in plant nutrient acquisition may vary with the season. Surprisingly, Furtado et al. (2019a) and Maciá-Vicente et al. (2016) did not observe seasonal changes in fungal communities colonizing *Salicornia europaea* and *S. patula*. However, we considered this surprising and aimed to confirm whether differences in host ploidy are potentially playing a role in seasonal variation.

Using a high-throughput amplicon sequencing technique, we investigated whether cytotype-specific fungal microbiomes occur in *Salicornia*. For this purpose, individuals of *S. europaea* (2x) and *S. procumbens* (4x) occurring in mixed-ploidy populations in the lower zone of a salt marsh located on the island of Spiekeroog (Germany) were collected during the spring and summer of 2020. We hypothesized that (*i) S. procumbens* (4x) has a more diverse root-associated endophytic fungal microbiome compared to *S. europaea* (2x), (*ii*) mutualistic fungi are more abundant in roots of *S. procumbens* (4x) than *S. europaea* (2x), and finally (iii) the observed patterns in microbiome composition are stable over two different sampling seasons. Additionally, we investigated whether *S. europaea* (2x) and *S. procumbens* (4x) show differences in root anatomical traits (e.g., cell size) which could impact differently fungal recruitment and microbiome assembly in the two cytotypes.

## Material and Methods

### Study area and sampling

Sampling was performed during spring (early June) and summer (late August) of 2020 in the lower zone of a salt marsh located within the limits of the Wadden Sea National Park on the island of Spiekeroog, Germany (53°45′N 7°43′E). Sampling was performed in the lower salt marsh (LSM) where *Salicornia europaea* (2x) and *S. procumbens* (4x) co-occur in mixed-ploidy populations. We focused our sampling on one population and one salt marsh zone in order to reduce potential cofounding effects of a larger scale spatial heterogenetiy, dispersal limitation and local adaptations on the structure of the fungal microbiomes associated to each cytotpe. To characterize the study area, the elemental chemical composition of a composite soil sample of each season (collected at three different points) was analysed (Table S1) at the Institute for Soil and Environment, Lufa Nord-West (Oldenburg, Germany).

We know from previous analyses (Buhk 2020) that position within the salt marsh has a significant influence on the root mycobiome of both *Salicornia* species. We, therefore, sampled both cytotypes in close proximity and overlapping root space. At the first sampling event (spring), 21 sampling points were randomly marked in the LSM (at least five meters apart from each other) and five individuals occurring side by side were collected at each point. Plants were carefully cleaned from the soil, large soil particles were removed from the root system and afterwards, each individual was placed in a separate plastic bag, labelled and kept at 4°C until further processing in the laboratory. In the summer, the sampling was performed identically, and individuals were collected at the same sampling points established in spring. In total, 100 individuals of unknown ploidy were collected in each season in order to maximize the chances of collecting a significant number of plants belonging to each cytotype.

In the laboratory, the roots and shoots of each individual were separated and processed separately. Shoots were washed, placed in plastic bags and kept at 4°C for a maximum of one week before genome size determination. Roots were washed to remove any adhering soil particle, cut in small pieces of about 1 cm and posteriorly, root pieces were surface sterilized following the protocol described in Crous et al. (2009). After that, the roots were kept at −20°C until DNA extraction.

### Genome size estimation and ploidy determination

For genome size estimation, the protocol described in Baranyi & Greilhuber (1996) was followed. Approximately 1 cm of shoots of sampled individuals was chopped with the same amount of an internal standard (*Hedychium gardnerianum;* 1C=2.01 pg; Meudt et al. 2015) into a homogenous mass by using a razor blade in a petri dish, containing 550 μl nuclei extraction buffer (OTTO I). After that, another 550 μl of the buffer was added to the sample suspension and filtered through a 30 μm CellTric filter (Partec GmbH, Münster, Germany) into a plastic tube, followed by the addition of 50 μl 5% RNAse. Samples were incubated in a water bath for 30 min at 37 °C. After that, 450μl of the cell suspension was transferred to a new plastic tube containing 2 mL of 6% propidium iodide staining solution. Staining was carried out in the darkness for one hour at 4°C. Genome size was determined in a CyFlow SL flow cytometer (Partec GmbH, Münster, Germany) equipped with a green laser (532 nm, 30 mW) as an excitation light source. Five thousand particles were studied and only measurements with coefficients of variation (CVs) < 5% were considered. However, for some individuals, measurements with CVs of 5-8% were also included. Ploidy measurements were performed until at least 20 individuals of each ploidy in each season was obtained. The distribution of each cytotype according to the different sampling points is shown in Table S2.

### Root anatomical investigation

In order to investigate whether *S. europaea* (2x) and *S. procumbens* (4x) show differences in root anatomical traits, additional samples of each cytotype were collected in the LSM and the ploidy level was estimated for species identification (Table S3). Root samples were washed under tap water and stored individually in ethanol (70%). For each cytotype, 10 individuals were selected and two root cuts of the primary root of each individual were obtained and used to measure the following anatomical parameters: maximum root diameter, periderma thickness, parenchyma thickness, diameter of the vascular cylinder and maximum diameter of parenchyma cells. Microscope slides were prepared, and measurements were performed using the software Olympus CellSens Entry 2.2 (Olympus Cooperation, Tokyo, Japan) under a Reichert-Jung Polyvar (Reichert, Wien, Austria) transmitted light microscope. For statistical analysis, a one-way ANOVA was performed, followed by a Tukey test to identify differences among cytotypes. Statistical analyses were performed in R (version 3.6.3, R Development Core Team, 2011).

### DNA extraction, PCR, library preparation and sequencing

DNA was extracted from 40 composite root samples (ten of each cytotype collected in each season). Roots of two individuals of the same ploidy that occurred in the same sampling point were grouped to form a composite sample for DNA extraction. In the case that only one diploid or tetraploid *Salicornia* was detected in a specific sampling point, a composite sample was formed with two samples of the same ploidy collected in different sampling points. This was performed due to the low root biomass content of some samples, especially those collected in spring. Before DNA extraction, root samples (approximately 20 mg) were lyophilized in an Alpha 1-2 LDplus freeze-dryer (Martin Christ Gefriertrocknungsanlagen GmbH, Germany) for 24 hours. The lyophilized samples were powdered using a Mixer Mill (Retsch, Germany) for two cycles of 1 min each with 30 oscillations per second. Genomic DNA was extracted using the innuPREP Plant DNA Kit (Analytic Jena AG, Jena, Germany), following the manufacturer’s instructions. The concentration and quality of extracted DNA were checked by absorption spectrophotometry with a Tecan Infinity 200 PRO microplate reader (Tecan Group Ltd., Männedorf, Switzerland). Dilutions for each sample were prepared to achieve a final concentration of 2 ng μl^-1^.

For the amplification of the ITS1 region of nrDNA, the primers ITS1FKyo2 (Toju et al. 2012) and ITS86R (Vancov & Keen 2009), which target all major fungal groups, were used. Each sample was amplified in duplicates and the reaction was prepared in a volume of 25 μl containing: 1.0 μl of DNA (2 ng μl^-1^), 5 μl of 5x Phusion Buffer, 0.6 μl of MgCl^2^, 0.5 μl 10 mM dNTPs, 0.5 μl of each primer (10 pmol μl^-1^), 0.2 μl of Phusion Hot Start Flex DNA polymerase (New England Biolabs GmbH) and 16.7 μl of nuclease-free water. PCR parameters were based on Maciá-Vicente et al. (2020) with minor adaptations: an initial step of 98°C for 1 min and 3 cycles of 98°C for 30 s, 55°C for 30 s and 72°C for 30 s. The further 32 cycles consisted of 98°C for 30 s, 55°C for 30 s, and 72°C for 30 s. The elongation step was at 68°C for 10 min. After amplification, products of the two reactions were pooled together and used in a second PCR reaction to ligate the adapters. This reaction was performed accordingly: an initial step of 98°C for 1 min and 7 cycles of 98°C for 30 s, 55°C for 30 s and 72°C for 30 s followed by an elongation step at 68°C for 10 min. PCR products were checked on a 1.2% agarose gel stained with ethidium bromide (Sigma-Aldrich). Subsequently, amplicons were sequenced at LGC Genomics (Berlin, Germany) in an Illumina MiSeq platform (V3 chemistry, 300□bp paired□end reads).

### Bioinformatics and statistical analyses

Sequence data was analysed and visualized using QIIME2 v.2020.11 (Bolyen et al. 2019). Demultiplexed sequences were processed using ITSxpress plugin (Rivers et al. 2018), which implements ITSx and BBMerge (Bushnell et al. 2017). Default parameters were used to run ITSxpress. Afterwards, reads were further processed using DADA2 denoise.paired plugin to identify exact amplicon sequence variants (ASVs) rather than lumping sequence variants to OTU’s (Callahan et al. 2017). Taxonomic assignments were performed using qiime feature-classifier classify-skylearn with a pre-trained Naïve-Bayes classifier using the database UNITE (version v.8.2) (Abarenkov et al. 2020). Amplicon sequence variants that were assigned to plants and protists were removed from the final ASV table. For functional assignment, a table containing the genera occurring in each cytotype was used and compared with the information available in the FungalTraits (Põlme et al. 2021) and FunGuild (Nguyen et al. 2016) databases.

In order to confirm whether sampling depth was sufficient, rarefaction curves were generated in QIIME2 using the qiime.diversity.alpha-rarefaction plugin. Differences in sampling depth were accounted by rarefying samples to a sampling depth of 1000 sequences before proceeding with diversity analyses. Alpha diversity was estimated based on Shannon diversity, observed richness, Faith’s phylogenetic diversity and Pielou’s evenness using a Kruskal-Wallis test. For beta diversity, a permutational analysis of variance (Adonis) was performed in order to check the significance of the factors ploidy and season, as well as their interaction. In addition, a Permanova test using 999 permutations was performed, and dissimilarities in microbial communities of *S. europaea* (2x) and *S. procumbens* (4x) in each sampling season was checked based on Bray-Curtis and Jaccard distances. Principal coordinate analysis (PCoA) plots were visualized in Emperor (Vázquez-Baeza et al. 2013). To define the core microbiome, we considered the definition of Astudillo-García et al. (2017), which defines as part of the core the ASVs present in at least 95% of samples with a minimum 1% relative abundance.

The files generated in QIIME2 were imported in R (version 3.6.3, R Development Core Team, 2011) using the package Qiime2R (https://github.com/jbisanz/qiime2R) and figures were generated using the package ggplot2 (Wickham 2016).

## Results

### Genome size estimation and ploidy

In total, genome size and ploidy of 135 individuals of *Salicornia* was estimated, resulting in 51 diploids (23 collected in spring and 28 in summer) and 84 tetraploids (45 collected in spring and 39 in summer). The mean genome size (2*C* DNA content) ± standard deviation was 0.652 pg ± 0.062 for *S. europaea* (2x) and 1.314 pg ± 0.061 for *S. procumbens* (4x).

### Root anatomical investigation

Significant differences between cytotypes were observed for the following characters: periderma thickness, parenchyma thickness and diameter of vascular cylinder (Table 1). *Salicornia procumbens* (4x) showed thicker periderma and parenchyma layers compared to *S. europaea* (2x), whereas *S. europaea* (2x) had a larger central cylinder compared to *S. procumbens* (4x). The presence of aerenchyma was observed in both *S. europaea* (2x) and *S. procumbens* (4x) (Figure 1).

**Figure 1.**
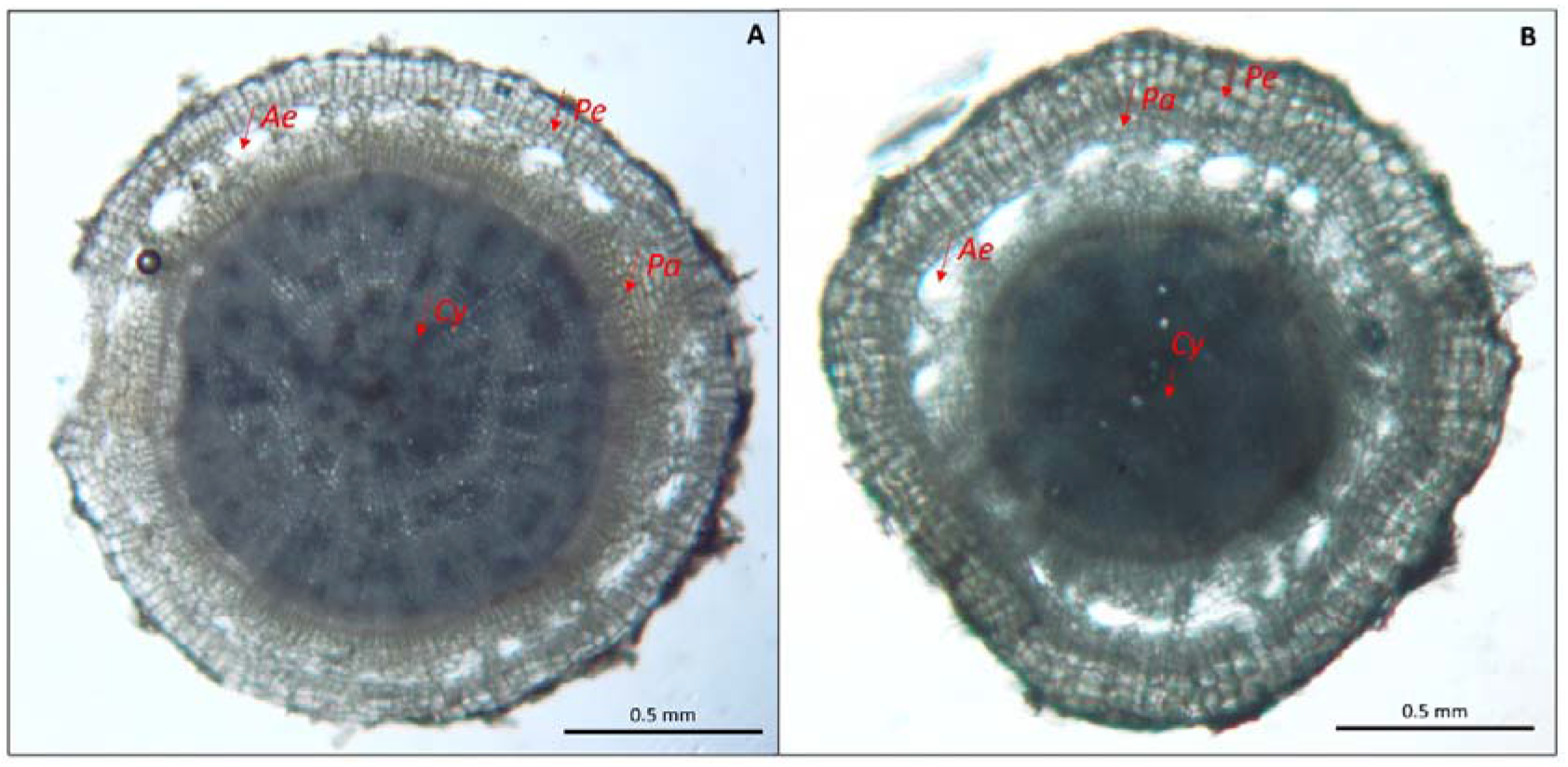
Root cross-section of *S. europaea* (2x) (A) and *S. procumbens* (4x) (B). *Pe:* periderma; *Pa*: parenchyma; *Ae*: Aerenchyma formation; *Cy*: central cylinder.

**Table 1.**
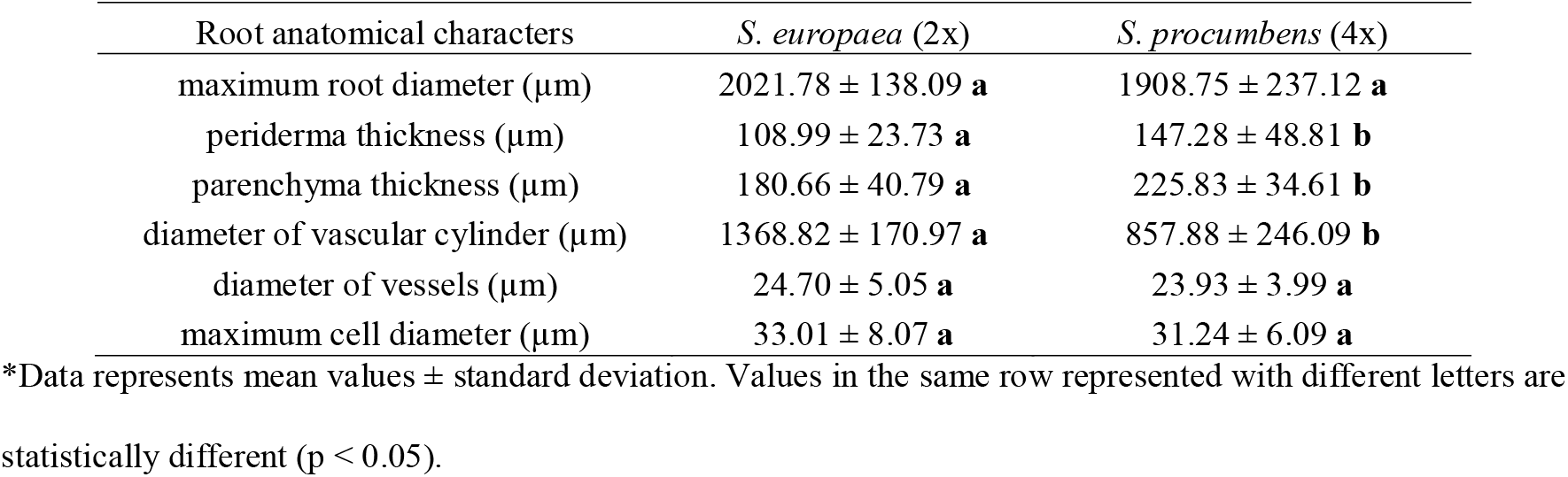
Root anatomical characters observed in *S. europaea* (2x) and *S. procumbens* (4x).

### Sequencing results

We initially obtained 1,284,546 sequences. After filtering out the sequences belonging to plants (87.83%) and Cercozoa (0.71%), a total of 145,546 high-quality sequences representing 122 exact amplicon sequence variants (ASVs) were obtained after denoising, merging, and chimera checking. The number of reads per sample ranged from 481 to 13431 with an average of 3093 sequences for *S. procumbens* (4x) and 5703 sequences for *S. europaea* (2x) samples. We rarefied the number of reads to 1000 reads per sample. After rarefaction, we remained with eight samples of *S. europaea* (2x) and five samples of. *S. procumbens* (4x) collected in spring; and ten samples of *S. europaea* (2x) and seven samples of *S. procumbens* (4x) collected in summer.

### Alpha and beta diversity of fungal communities

Shannon diversity-based rarefaction curves for both cytotypes showed a plateau at 200 sequences (Figure S1), indicating that sampling depth was enough to cover the diversity of the samples. A similar tendency was observed for richness-based rarefaction curves (Figure S2). Alpha diversity analyses showed higher Shannon Diversity (p=0.012) and Pielou’s Evenness (p=0.019) indices in *S. procumbens* (4x) samples compared to *S. europaea* (2x) collected in spring (Figure 2). On the other hand, no significant differences but strong trends in the same direction were observed for samples collected in summer (Shannon Diversity Index, p=0.118; Pielou’s Evenness, p=0.078). No significant differences were observed in Faith’s Phylogenetic Diversity in spring (p=0.417) and summer (p=0.625) and observed richness in spring (p=0.077) and summer (p=0.116) (Figure 2). When the effect of season was analyzed separately for each cytotype, no differences were observed in any of the measured alpha-diversity indices (Table S4).

**Figure 2.**
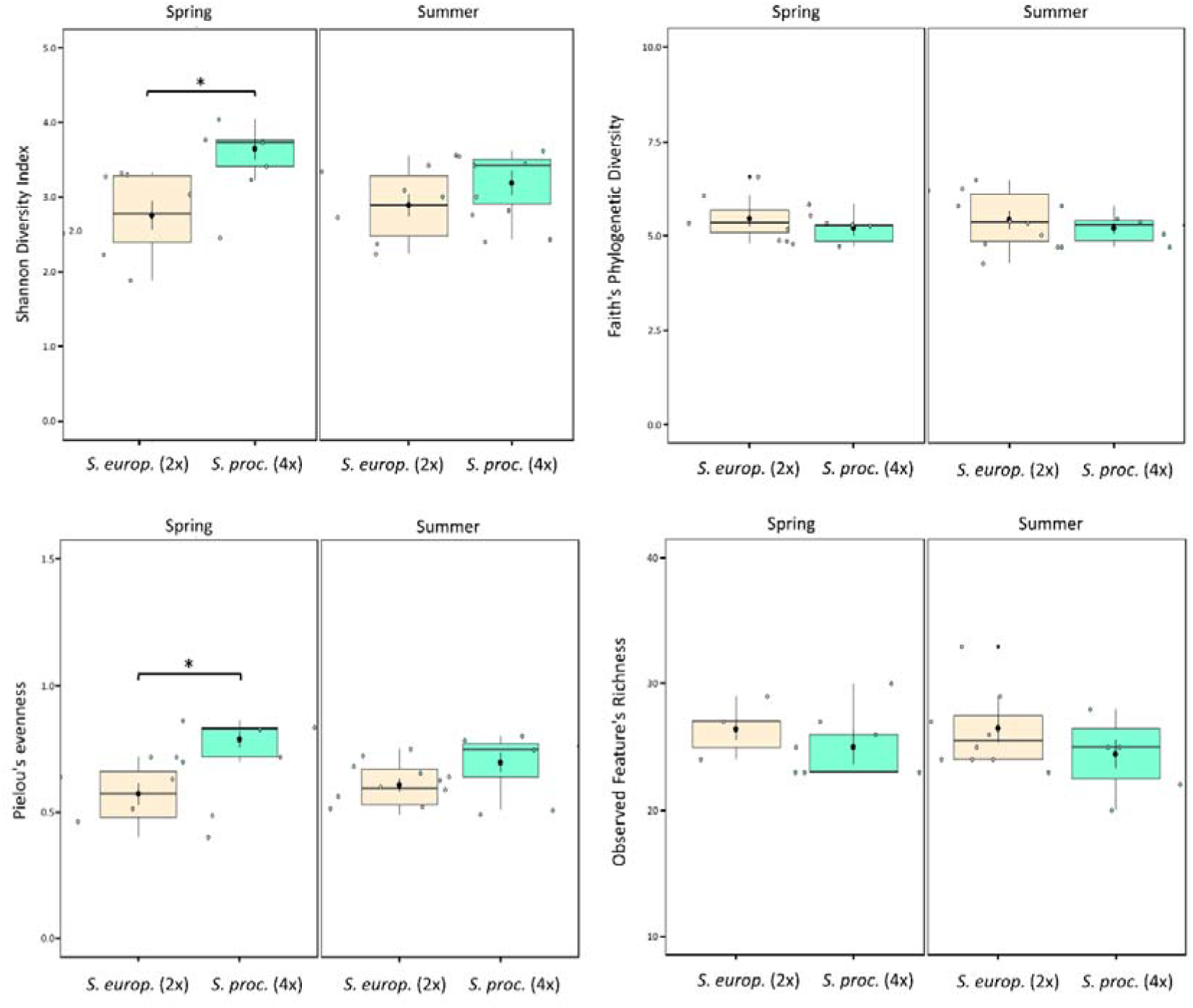
Boxplots of Shannon diversity, Faith’s phylogenetic diversity, Pielou’s evenness and observed richness for *S. europaea* (2x) and *S. procumbens* (4x) in spring and summer. Significant differences (Kruskal-Wallis test, P-value < 0.05) are represented by *.

Beta diversity analyses showed significant differences in fungal community between the two cytotypes (ADONIS, p=0.014), whereas season (p=0.246) and the interaction between these factors were not statistically significant (p=0.604) (Table 2). In spring, no differences were observed between *S. europaea* (2x) and *S. procumbens* (4x) based on Bray-Curtis (Permanova, p=0.331, Figure 3A) and Jaccard distances (Permanova, p=0.476, Figure 3C); however, in summer, samples of *S. procumbens* (4x) clustered together and significantly differed from that of *S. europaea* (2x) based on both Bray-Curtis (Permanova, p=0.04, Figure 3B) and Jaccard distances (Permanova, p=0.009, Figure 3D).

**Figure 3.**
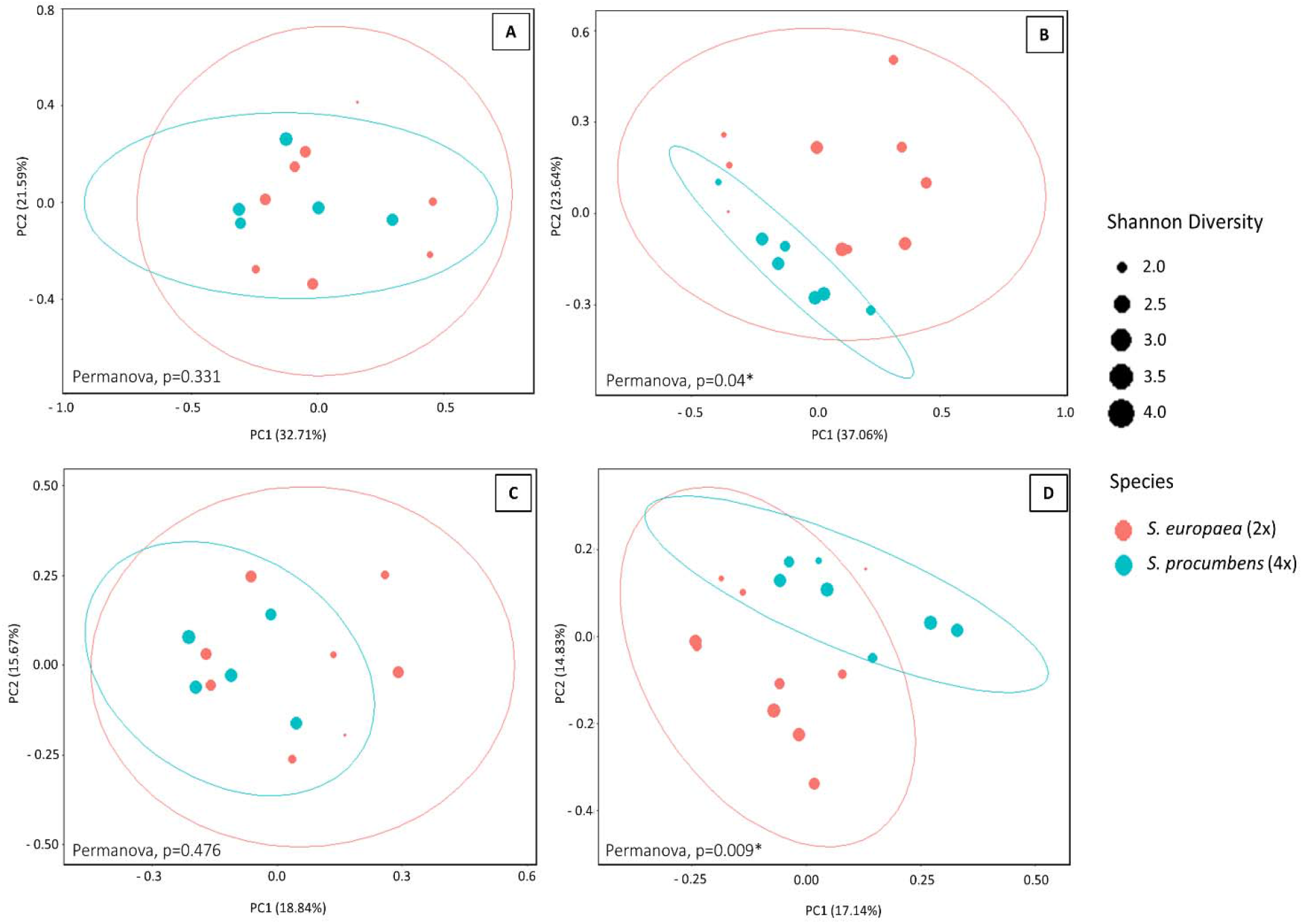
Principal Coordinate Analysis (PCoA) based on Bray-Curtis distances showing the distribution of fungal communities of *S. europaea* (2x) and *S. procumbens* (4x) in spring (**A**) and summer (**B**). PCoA based on Jaccard distances in spring (**C**) and summer (**D**). Variation explained by the data in PCoA is presented in percentage.

**Table 2.** Permutational analysis of variance (ADONIS) for the factors season and ploidy.

|  | D<sub>f</sub> | Sum Sq. | Mean Sq. | F model | R<sup>2</sup> | P-value |
| --- | --- | --- | --- | --- | --- | --- |
| Season | 1 | 0.251 | 0.251 | 1.296 | 0.042 | 0.246 |
| Ploidy | 1 | 0.529 | 0.529 | 2.731 | 0.088 | 0.014 |
| Season:Ploidy | 1 | 0.151 | 0.151 | 0.779 | 0.025 | 0.604 |
| Residuals | 33 | 5.045 | 0.194 |  | 0.843 |  |
| Total | 36 | 5.977 |  |  | 1 |  |

### Core microbiome

We defined the core microbiome as ASVs present in at least 95% of samples with minimum 1% relative abundance in both *S. europaea* (2x) and *S. procumbens* (4x) samples. Overall, 13 ASVs were identified: ten belonging to Ascomycota (unidentified *Xylariales, Laburnicola* sp., *Lentithecium pseudoclioninum, Cladosporium* sp., *Neocamarosporium* sp., *Alternaria dactylidicola*, unidentified *Hypocrales, Fusarium* sp., *Plectosphaerella* sp., *Alternaria chlamydospora*) and three to Basidiomycota (unidentified *Malasseziales, Wallemia* sp., *Malassezia restricta*). However, there was no core microbiome taxon associated with a single cytotype based on this definition.

### Taxonomical composition and functional assignment

Fungi belonging to three phyla were detected in the roots of *S. europaea* (2x) (*Ascomycota* 70.2%, *Basidiomycota* 29.1% and *Chytridiomycota* 0.7%) and *S. procumbens (Ascomycota* 60.5%, *Basidiomycota* 36.9% and *Chytridiomycota* 2.6%). The orders *Xylariales* and *Pleosporales* (Ascomycota), where many DSE fungi are classified, along with *Malasseziales* (Basidiomycota) corresponded to 85% of the sequences in *S. europaea* and 75 % in *S. procumbens* (4x) (Figure 4A). Figure 4B shows the taxon bar plots of fungal genera associated with each cytotype in each season. Whereas 25 genera were found to be associated with both cytotypes, six genera were exclusively associated with *S. procumbens* (4x) (Ascomycota: *Lasiodiplodia, Didymella, Ophiosphaerella* and Basidiomycota: *Cystofilobasidium, Tausonia, Solicoccozyma*) and ten with *S. europaea* (2x) (Ascomycota: *Neodidymelliopsis, Preussia, Trematosphaeria, Acremonium* and Basidiomycota: *Kurtzmanomyces, Microstroma, Rhodotorula, Sporobolomyces, Itersonilia* and *Papiliotrema*) (Figure 4D). Typical genera of DSE were detected, for example, the genera *Alternaria, Cladosporium, Ophiosphaerella* and *Paraphaeosphaeria*. The majority of the genera were functionally assigned to saprophytic or plant pathogenic trophic modes (Table S5). The most frequent taxon for which it was possible to assign species identity was *Lentithecium pseudoclioninum* (Ascomycota, Pleosporales).

**Figure 4.**
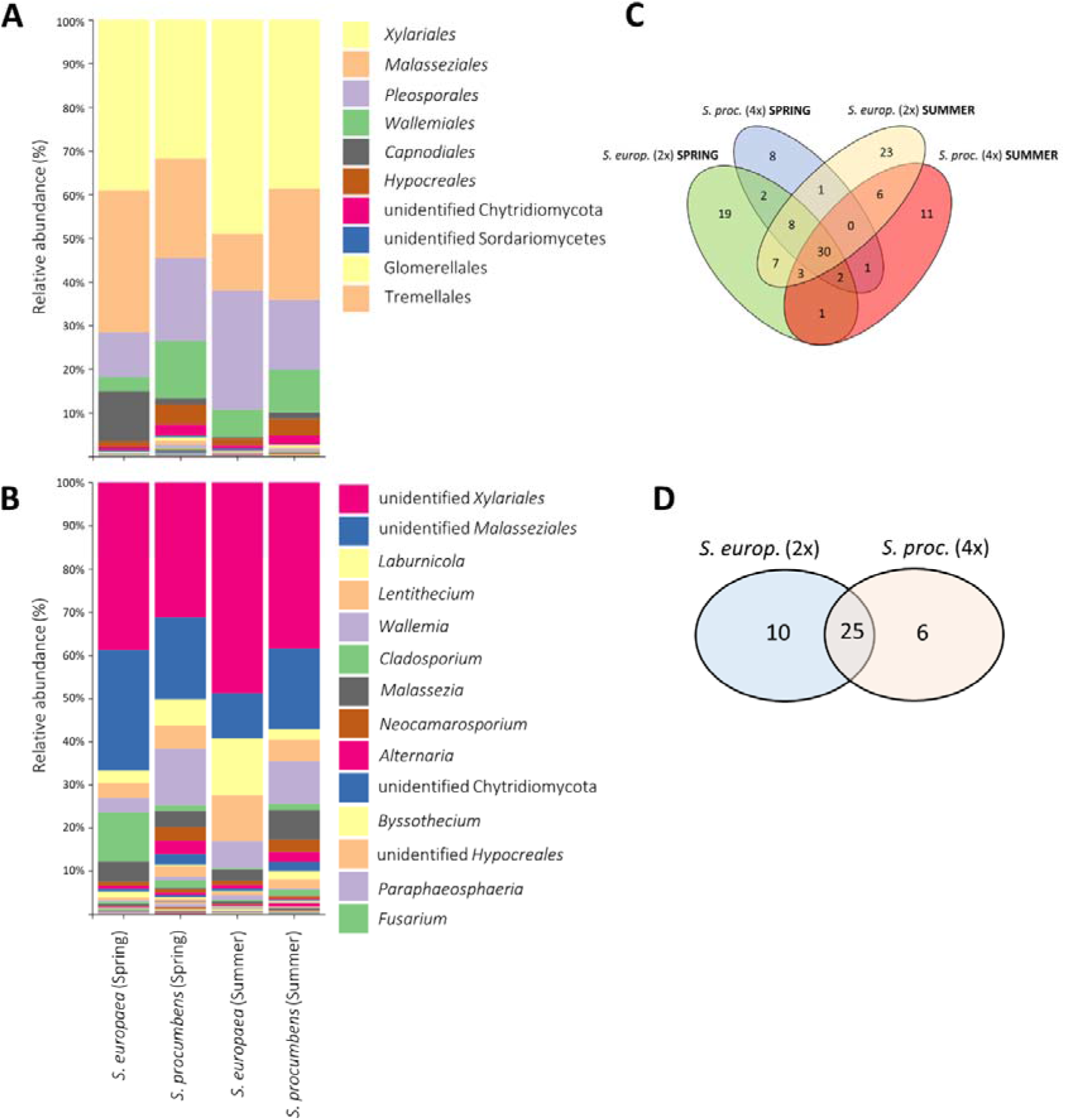
Fungal community composition of *S. europaea* (2x) and *S. procumbens* (4x). **(A)** Relative read abundance of fungal orders present in each cytotype collected in spring and summer. **(B)** Relative read abundance of fungal genera present in each cytotype collected in spring and summer. **(C)** Venn diagram of ASVs associated with each cytotype collected in each season. **(D)** Venn diagram of fungal genera associated with each cytotype.

## Discussion

Polyploidy induces diverse genetic and phenotypic changes in plants. Consequently, there can be cascading effects in their interaction with microorganisms (Segraves & Anneberg 2016), which potentially allows differentiation of polyploids from their diploid relatives. We hypothesized that *S. procumbens* (4x) harbor more diverse fungal communities compared to *S. europaea* (2x) based on greater physiological demands (e.g., higher photosynthetic rates and nutrient requirements) and anatomical differences (e.g., larger cell size) observed in polyploids (Forrester & Ashman 2018; Šmarda et al. 2013). Higher physiological demands may require *S. procumbens* (4x) to establish association with a broader range of beneficial fungal partners whereas larger cells provide more space for different fungi to colonize. Our results indicate significant differences in the root-endophytic fungal microbiome associated with *S. procumbens* (4x) and *S. europaea* (2x) in terms of alpha diversity, with *S. procumbens* (4x) samples showing higher Shannon diversity and evenness indices but a trend to lower richness (Fig. 2). This pattern is more prominent in spring suggesting that the fungal community settles by several spring colonizers lost in the year and only few fungi colonizing the plants later. Further, beta diversity analysis showed dissimilar fungal communities associated with *S procumbens* (4x) and *S. europaea* (2x) samples collected in summer (Fig. 3). However, when the effect of seasonality was independently analyzed, no differences were observed between spring and summer, which indicates comparatively low turnover during the season. The same was reported by Furtado et al. (2019a) when fungal communities colonizing *S. europaea* in spring and autumn were analysed. Finally, we did not observe larger cells in *S. procumbens* (4x) but a thicker parenchyma layer, which suggests a larger area for fungal colonization.

We investigated cytotype-specific root anatomical traits since previous studies have shown that they can affect microbial recruitment and community assembly. For example, Forrester et al. (2020) noted a greater generalization in bacterial assemblages in autotetraploid individuals of *Medicago sativa* compared to diploids, suggesting that larger cells in polyploids could host a greater quantity of different bacterial symbionts, compared to smaller cells in diploids. In the case of fungi, Galindo-Castañeda et al. (2019) reported a decrease in mycorrhizal colonization in *Zea mays* with a decrease in the living cortical area. In the case of *Salicornia*, our anatomical data showed no differences in cell size between the two cytotypes (Table 1). However, *S. europaea* (2x) showed a larger central cylinder compared to *S. procumbens* (4x) whereas thicker periderma and parenchyma layers were observed in *S. procumbens* (4x) along with an abundant ocurrence of aerenchyma (Fig. 1, Table 1). We speculate that a thicker periderma and parenchyma, along with the presence of air spaces in *S. procumbens* (4x) roots, could provide advantages for endophytes by providing space and oxygen for fungal establishment and survival, especially when the anoxic conditions of salt marshes are considered. Another point to be further investigated is whether the abundant occurrence of aerenchyma, especially in *S. procumbens* (4x), could be due to the presence of specific fungal symbionts. The induction of aerenchyma by fungi was reported by Hu et al. (2018), who observed that the inoculation of rice plants with *Phomopsis liquidambari* (Ascomycota) directly influenced aerenchyma formation via the accumulation of indole-3-acetic acid and ethylene. A similar pattern could also be occurring with *S. procumbens* (4x) but with different fungal symbionts. Although not investigated in this study, host physiology could also be affecting fungal recruitment. Micallef et al. (2019) demonstrated that exudation patterns differed between genotypes of *Arabidopsis thaliana* affecting bacterial recruitment. The same was observed by Zhalnina et al. (2018) in different genotypes of *Avena barbata*. Similar processes are likely to occur with fungi. Since whole-genome duplication can alter root exudation patterns (Jesus-Gonzalez & Weathers 2003), it is possible that in *Salicornia* as well, the two cytotypes produce different root exudates, which consequently leads to different fungal recruitment and community establishment.

The results of alpha diversity analyses (Fig. 2) showed more diverse and even fungal communities associated with *S. procumbens* (4x) compared to *S. europaea* (2x), although significant differences were observed only in spring. The lower Shannon diversity and evenness observed in *S. europaea* (2x) is seemingly due to the presence of a few and highly dominant fungal taxa colonizing the roots of this cytotype (e.g. unidentified *Xylariales* and *Malasseziales, Cladosporium* sp.) but many rare species. This pattern has also been observed in the root-associated bacterial and fungal microbiomes of other plant species (Diaz-Garza et al. 2020, Gobbi et al. 2020). Two possible reasons for this are either, highly abundant fungi in *S. europaea* (2x) compact the remaining fungal assemblage, outcompeting other species and reducing community evenness. Alternatively, the roots of the diploid species form a specialized environment allowing only a few species to establish. Beta diversity analyses showed that fungal communities clustered according to cytotype and did not cluster according to season (Table 2). Other studies investigating the influence of seasonality in shaping root-endophytic fungal microbiomes in *Salicornia* also showed that community composition and structure remained stable over the complete *Salicornia patula* lifecyle (Maciá-Vicente et al. 2016) and in the case of *S. europaea*, it did not change between spring and autumn (Furtado et al. 2019a). Thus, despite being annual these plants seem not to be limited by a lack of mutualistic endophytic fungi in the early season but are also immediately affected by pathogenic fungi.

The core microbiome of *Salicornia* comprised 13 ASVs. Some of these taxa have been already reported colonizing *Salicornia* spp. such as *Alternaria chlamydospora* (Furtado et al. 2019a), *Cladosporium* sp. (Furtado et al. 2019b; Maciá-Vicente et al. 2016) and *Fusarium* sp. (Furtado et al. 2019b). In addition, other taxa detected in our study have been reported colonizing other chenopod hosts, such as *Laburnicola* sp. isolated from *Suaeda salsa* (Yuan et al. 2020) and *Neocamarosporium* sp. isolated from *Atriplex portulacoides* (Gonçalves et al. 2019). We detected some ASVs exclusively associated with each cytotype. For example, an unidentified ASV belonging to Tremellomycetes was detected in five samples of *S. europaea* (2x). The same was observed for a unidentified ASV belonging to Pleosporales in *S. procumbens* (4x). Nevertheless, these do not constitute cytotype-specific core microbiomes since these ASVs were not detected in more than 95% of the samples of each cytotype. Apart from the genera *Alternaria* sp. and *Fusarium* sp., well known as plant pathogens but also reported as mutualists (Mendoza & Siroka 2009; Musetti et al. 2007) and the genus *Laburnicola* sp., reported in a mutualistic association with the halophyte *Suaeda salsa* (Yuan et al. 2020), the other taxa present in the *Salicornia* core microbiome are mostly considered saprotrophs (e.g, order *Xylariales, Cladosporium* sp., *Lentithecium* sp.) or plant pathogens (e.g., *Cladosporium* sp.).

As observed for the core microbiome, functional assignment revealed that most of the genera detected in our samples were classified in two trophic modes: saprotroph or pathotroph. However, two genera (*Alternaria* and *Fusarium*) colonizing both *S. europaea* (2x) and *S. procumbens* (4x), and *Acremonium*, detected in roots of only *S. europaea* (2x), were classified into the pathotroph-saprotroph-symbiotroph trophic mode, according to FunGuild database (Nguyen et al. 2016). The outcome of the plant-fungal association is known to depend on a set of abiotic and biotic factors (Hardoim et al. 2015). Studies have shown that fungi may change lifestyle (e.g., from mutualistic to pathogenic) depending on host, plant genotype, environmental conditions and the dynamic network of interactions within the plant microbiome (Fesel & Zuccaro 2016; Hardoim et al. 2015), whereas the effect of whole-genome duplication in this regard is still unknown. Physiological characteristics inherent to each host, for example, the differences in fungal gene expression in response to the host or differences in plant’s ability to respond to the fungus, has been shown to play a major role in controlling the outcome of the symbiosis (Redman et al. 2001). Interestingly, plant physiology is known to differ between diploids and polyploids (Baker et al. 2017), although not yet explored in *Salicornia*. It is possible that cytotype-specific physiological characteristics in *Salicornia*, such as different nutrient requirements, could modulate in different ways the symbiotic interaction with endophytic fungi.

We hypothesized that in the salt marsh context, *S. procumbens* (4x) has a larger microscale distribution, possibly due to its association with more or more generalized mutualistic fungi. However, there are no studies to investigate whether establishing symbiosis with beneficial fungi is also part of the strategy adopted by polyploids to colonize different habitats. In the case of *Salicornia*, Teege et al. (2011) detected an ecologically differentiated distribution of two closely-related *Salicornia* species in the salt marsh zones suggesting a strong selection during seed germination and seedling establishment as reasons for the habitat differentiation. Nevertheless, there is no information regarding the influence of belowground interactions in this regard. Since polyploids have higher demands for nutrients in order to synthesize additional chromosomal sets (Guignard et al. 2016; Šmarda et al. 2013), associating with mutualistic fungi could be part of their strategy to fulfil the nutrient requirements and increase its competitiveness. Our results showed that *S. procumbens* (4x) did not associate specifically with mutualistic fungal symbionts. This finding leaves the possibility that either the assignments to guilds are inexact because of the high functional differences at species or genus level or fungal endophytes are not contributing or are playing a minor role in *S. procumbens* (4x) colonization of harsh habitats, such as tidal flats where frequent flooding associated to high salinity occur. However, we cannot exclude that association with mutualistic fungi is habitat-dependent. Therefore, in a next step, sampling of *Salicornia* across the salt marsh into the pioneer zone and tidal flats would be necessary to understand whether mutualists are more abundant in roots of tetraploid plants colonizing harsher parts of the salt marsh and not the LSM.

Overall, our results corroborate past work in which polyploid plants were found to harbor more diverse root-associated endophytic communities compared to diploids, not only fungi (Tĕšitelová et al. 2013) but also bacteria (Wipf & Coleman-Derr 2021). However, further research should consider including samples of *S. europaea* (2x) and *S. procumbens* (4x) collected in different populations. This would allow for drawing conclusions in a broader ecological context. In addition to that, further investigations are necessary to elucidate the exact anatomical or physiological mechanisms inherent to each cytotype and responsible for these differences. It is possible that larger parenchyma observed in *S. procumbens* (4x) could provide more space for fungal colonization also benefiting nutrient uptake. To support this hypothesis, further work under controlled conditions is necessary to investigate whether *S. procumbens* (4x) has different physiological demands (e.g., higher nutrient requirements) compared to *S. europaea* (2x). Finally, since DNA-based methods do not provide information on the active root-inhabiting fungi colonizing each cytotype, transcriptomic analyes would help us to understand whether fungal communities are dominated in their activity by few species and whether different active fungal community profiles occur in the two cytotypes.

## Declarations

### Availability of data and material

All raw sequences are available from BioProject ID: PRJNA764035.

### Author contributions

DRG and DCA planned and designed the study with suggestions provided by RP. DRG performed sampling, molecular work, data analyses and wrote the first draft. DRG, RP and DCA contributed to the final version of the manuscript.

## Acknowledgements

We would like to thank Sabrina Schöngart for performing the ploidy measurements and Eike Mayland-Quellhorst for his help with molecular work. We also thank the administration of the Lower Saxony Wadden Sea National Park for allowing us to collect the samples used in this study.

