## Supplementary material for "Polyploidy and plant-fungus symbiosis: evidence of cytotype-specific microbiomes in the halophyte *Salicornia* (Amaranthaceae)"

**Table S1:** Elemental chemical composition of a composite soil sample collected in the lower salt marsh in Spring and Summer. EC: electrical conductivity. Values are means of four replicates and standard deviation.

|  | **pH (CaCl_2_)** | **EC (mS/cm)** | **Nitrogen (%)** | **Carbon (%)** | **C/N Ratio** | **P_2_O_5_ (mg/100g)** | **K_2_O (mg/100g)** | **Mg (mg/100g)** |
| --- | --- | --- | --- | --- | --- | --- | --- | --- |
| ***Spring*** | 6.97 ± 0.05 | 11.35 ± 0.49 | 0.57 ± 0.04 | 6.27 ± 0.45 | 11.00 ± 0.05 | 27.33 ± 1.70 | 162.67 ± 14.08 | 232.33 ± 45.07 |
| ***Summer*** | 6.91 ± 0.06 | 12.14 ± 0.56 | 0.52 ± 0.08 | 4.86 ± 0.94 | 9.33 ± 0.88 | 7.77 ± 2.08 | 116.33 ± 7.52 | 218.17 ± 3.93 |

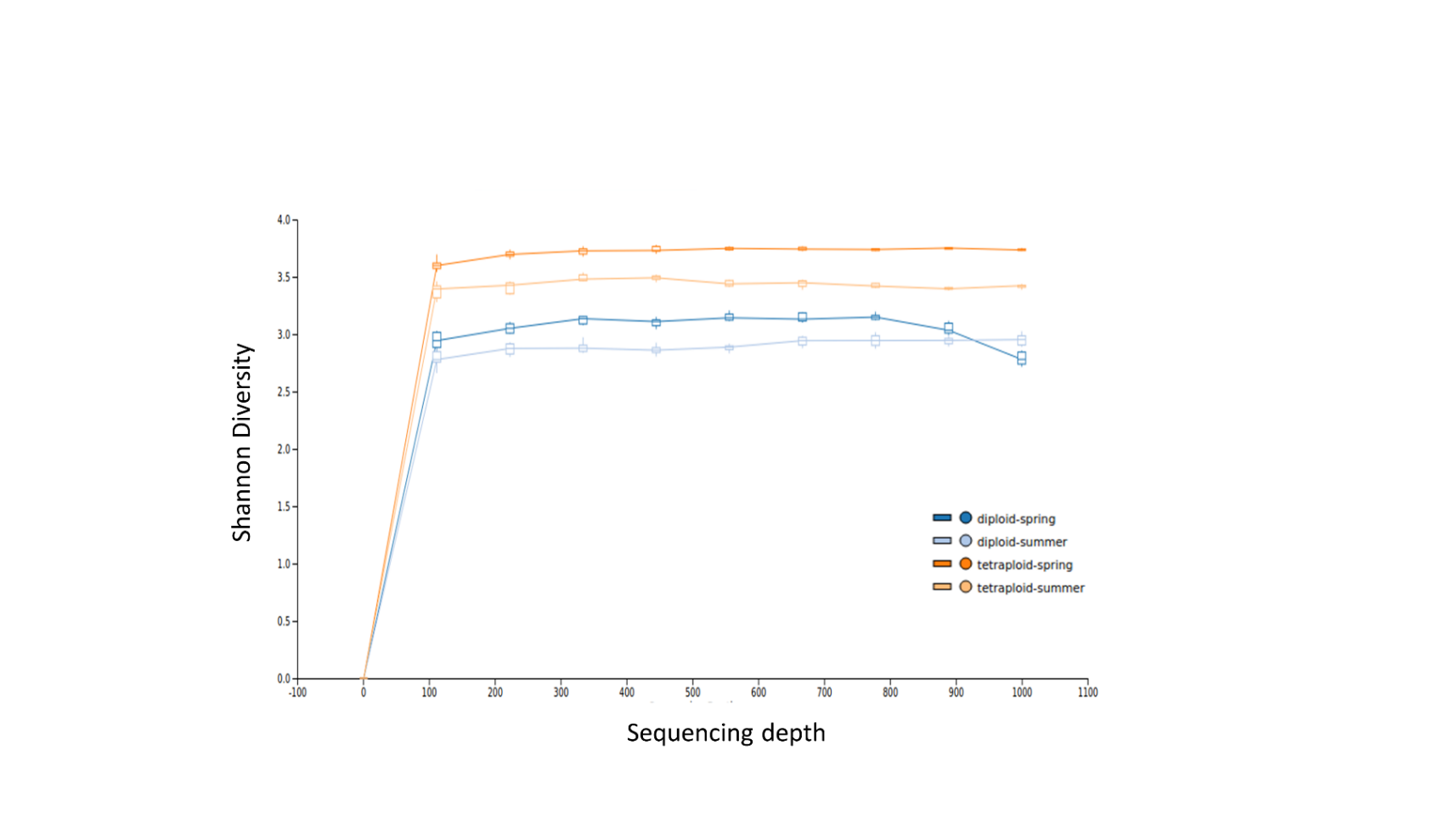

**Figure S1.** Shannon diversity-based rarefaction curves.

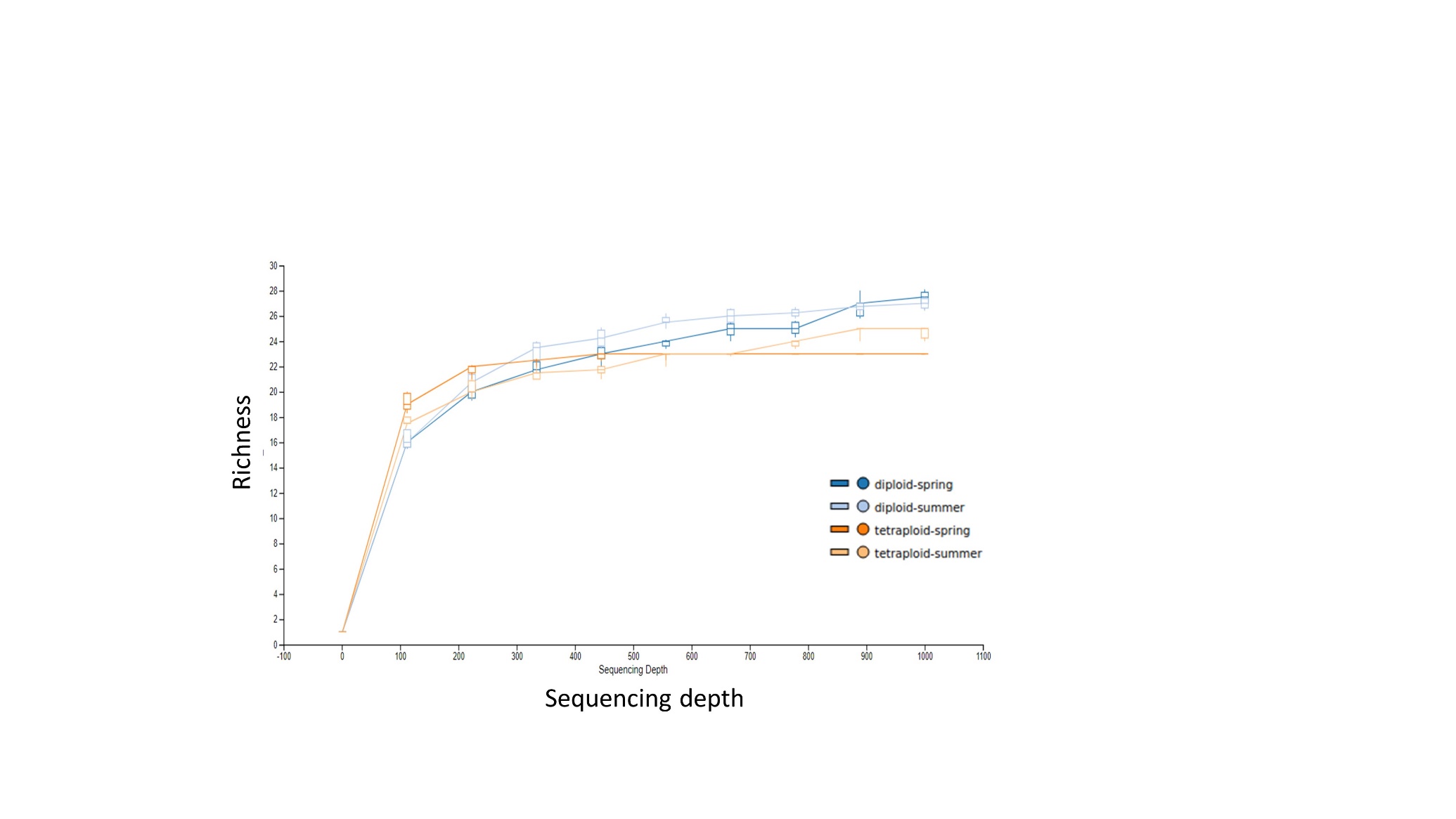

**Figure S2.** Richness-based rarefaction curves.

**Table S2**. Distribution of *S. europaea* and *S. procumbens* collected in this study according to the different sampling points. Samples presented in this table were used for molecular analyses.

| **Sample ID** | **Month** | **Zone** | **Point** | **Genome Size (2C DNA content, pg)** | **Ploidy** | **Species** |
| --- | --- | --- | --- | --- | --- | --- |
| DG-Sa-P1.1 | June | LSM | 1 | 1.320 | tetraploid | *Salicornia procumbens* |
| DG-Sa-P1.2 | June | LSM | 1 | 0.646 | diploid | *Salicornia europaea* |
| DG-Sa-P1.3 | June | LSM | 1 | 0.614 | diploid | *Salicornia europaea* |
| DG-Sa-P1.4 | June | LSM | 1 | 0.634 | diploid | *Salicornia europaea* |
| DG-Sa-P1.5 | June | LSM | 1 | 0.700 | diploid | *Salicornia europaea* |
| DG-Sa-P2.1 | June | LSM | 2 | 1.271 | tetraploid | *Salicornia procumbens* |
| DG-Sa-P2.2 | June | LSM | 2 | 1.320 | tetraploid | *Salicornia procumbens* |
| DG-Sa-P2.3 | June | LSM | 2 | 1.318 | tetraploid | *Salicornia procumbens* |
| DG-Sa-P2.4 | June | LSM | 2 | 1.288 | tetraploid | *Salicornia procumbens* |
| DG-Sa-P2.5 | June | LSM | 2 | 0.644 | diploid | *Salicornia europaea* |
| DG-Sa-P3.1 | June | LSM | 3 | 1.278 | tetraploid | *Salicornia procumbens* |
| DG-Sa-P3.2 | June | LSM | 3 | 1.274 | tetraploid | *Salicornia procumbens* |
| DG-Sa-P3.3 | June | LSM | 3 | 1.263 | tetraploid | *Salicornia procumbens* |
| DG-Sa-P4.3 | June | LSM | 4 | 0.677 | diploid | *Salicornia europaea* |
| DG-Sa-P4.1 | June | LSM | 4 | 1.313 | tetraploid | *Salicornia procumbens* |
| DG-Sa-P4.2 | June | LSM | 4 | 1.293 | tetraploid | *Salicornia procumbens* |
| DG-Sa-P5.1 | June | LSM | 5 | 1.337 | tetraploid | *Salicornia procumbens* |
| DG-Sa-P5.2 | June | LSM | 5 | 0.644 | diploid | *Salicornia europaea* |
| DG-Sa-P5.3 | June | LSM | 5 | 1.305 | tetraploid | *Salicornia procumbens* |
| DG-Sa-P6.1 | June | LSM | 6 | 1.325 | tetraploid | *Salicornia procumbens* |
| DG-Sa-P6.2 | June | LSM | 6 | 1.393 | tetraploid | *Salicornia procumbens* |
| DG-Sa-P6.3 | June | LSM | 6 | 1.352 | tetraploid | *Salicornia procumbens* |
| DG-Sa-P7.2 | June | LSM | 7 | 0.683 | diploid | *Salicornia europaea* |
| DG-Sa-P7.1 | June | LSM | 7 | 1.322 | tetraploid | *Salicornia procumbens* |
| DG-Sa-P7.3 | June | LSM | 7 | 1.283 | tetraploid | *Salicornia procumbens* |
| DG-Sa-P8.1 | June | LSM | 8 | 0.644 | diploid | *Salicornia europaea* |
| DG-Sa-P8.2 | June | LSM | 8 | 0.631 | diploid | *Salicornia europaea* |
| DG-Sa-P8.3 | June | LSM | 8 | 1.255 | tetraploid | *Salicornia procumbens* |
| DG-Sa-P9.1 | June | LSM | 9 | 1.298 | tetraploid | *Salicornia procumbens* |
| DG-Sa-P9.2 | June | LSM | 9 | 1.282 | tetraploid | *Salicornia procumbens* |
| DG-Sa-P9.3 | June | LSM | 9 | 1.326 | tetraploid | *Salicornia procumbens* |
| DG-Sa-P10.1 | June | LSM | 10 | 1.298 | tetraploid | *Salicornia procumbens* |
| DG-Sa-P10.2 | June | LSM | 10 | 1.316 | tetraploid | *Salicornia procumbens* |
| DG-Sa-P10.3 | June | LSM | 10 | 1.313 | tetraploid | *Salicornia procumbens* |
| DG-Sa-P11.1 | June | LSM | 11 | 0.623 | diploid | *Salicornia europaea* |
| DG-Sa-P11.2 | June | LSM | 11 | 0.626 | diploid | *Salicornia europaea* |
| DG-Sa-P11.3 | June | LSM | 11 | 1.319 | tetraploid | *Salicornia procumbens* |
| DG-Sa-P11.4 | June | LSM | 11 | 0.622 | diploid | *Salicornia europaea* |
| DG-Sa-P11.5 | June | LSM | 11 | 0.643 | diploid | *Salicornia europaea* |
| DG-Sa-P12.1 | June | LSM | 12 | 0.653 | diploid | *Salicornia europaea* |
| DG-Sa-P12.2 | June | LSM | 12 | 1.275 | tetraploid | *Salicornia procumbens* |
| DG-Sa-P12.3 | June | LSM | 12 | 1.335 | tetraploid | *Salicornia procumbens* |
| DG-Sa-P13.1 | June | LSM | 13 | 1.348 | tetraploid | *Salicornia procumbens* |
| DG-Sa-P13.2 | June | LSM | 13 | 0.613 | diploid | *Salicornia europaea* |
| DG-Sa-P13.3 | June | LSM | 13 | 0.636 | diploid | *Salicornia europaea* |
| DG-Sa-P14.1 | June | LSM | 14 | 1.209 | tetraploid | *Salicornia procumbens* |
| DG-Sa-P14.2 | June | LSM | 14 | 1.265 | tetraploid | *Salicornia procumbens* |
| DG-Sa-P14.3 | June | LSM | 14 | 0.626 | diploid | *Salicornia europaea* |
| DG-Sa-P15.1 | June | LSM | 15 | 0.597 | diploid | *Salicornia europaea* |
| DG-Sa-P15.2 | June | LSM | 15 | 0.639 | diploid | *Salicornia europaea* |
| DG-Sa-P15.3 | June | LSM | 15 | 0.679 | diploid | *Salicornia europaea* |
| DG-Sa-P15.4 | June | LSM | 15 | 1.327 | tetraploid | *Salicornia procumbens* |
| DG-Sa-P15.5 | June | LSM | 15 | 0.641 | diploid | *Salicornia europaea* |
| DG-Sa-P16.1 | June | LSM | 16 | 1.268 | tetraploid | *Salicornia procumbens* |
| DG-Sa-P16.2 | June | LSM | 16 | 1.260 | tetraploid | *Salicornia procumbens* |
| DG-Sa-P17.1 | June | LSM | 17 | 1.381 | tetraploid | *Salicornia procumbens* |
| DG-Sa-P17.2 | June | LSM | 17 | 1.265 | tetraploid | *Salicornia procumbens* |
| DG-Sa-P18.1 | June | LSM | 18 | 1.349 | tetraploid | *Salicornia procumbens* |
| DG-Sa-P18.2 | June | LSM | 18 | 1.286 | tetraploid | *Salicornia procumbens* |
| DG-Sa-P19.1 | June | LSM | 19 | 1.358 | tetraploid | *Salicornia procumbens* |
| DG-Sa-P19.2 | June | LSM | 19 | 1.304 | tetraploid | *Salicornia procumbens* |
| DG-Sa-P20.1 | June | LSM | 20 | 1.414 | tetraploid | *Salicornia procumbens* |
| DG-Sa-P20.2 | June | LSM | 20 | 1.732 | tetraploid | *Salicornia procumbens* |
| DG-Sa-P21.1 | June | LSM | 21 | 0.596 | diploid | *Salicornia europaea* |
| DG-Sa-P21.2 | June | LSM | 21 | 1.296 | tetraploid | *Salicornia procumbens* |
| DG-Sa-P21.3 | June | LSM | 21 | 1.319 | tetraploid | *Salicornia procumbens* |
| DG-Sa-P21.4 | June | LSM | 21 | 1.284 | tetraploid | *Salicornia procumbens* |
| DG-Sa-P21.5 | June | LSM | 21 | 1.269 | tetraploid | *Salicornia procumbens* |
| DG-Sa-A-P1.1 | August | LSM | 1 | 0.616 | diploid | *Salicornia europaea* |
| DG-Sa-A-P1.2 | August | LSM | 1 | 0.542 | diploid | *Salicornia europaea* |
| DG-Sa-A-P1.3 | August | LSM | 1 | 0.609 | diploid | *Salicornia europaea* |
| DG-Sa-A-P1.4 | August | LSM | 1 | 0.631 | diploid | *Salicornia europaea* |
| DG-Sa-A-P1.5 | August | LSM | 1 | 0.968 | diploid | *Salicornia europaea* |
| DG-Sa-A-P1.6 | August | LSM | 1 | 0.639 | diploid | *Salicornia europaea* |
| DG-Sa-A-P2.1 | August | LSM | 2 | 1.258 | tetraploid | *Salicornia procumbens* |
| DG-Sa-A-P2.2 | August | LSM | 2 | 0.621 | diploid | *Salicornia europaea* |
| DG-Sa-A-P2.3 | August | LSM | 2 | 0.623 | diploid | *Salicornia europaea* |
| DG-Sa-A-P2.4 | August | LSM | 2 | 1.285 | tetraploid | *Salicornia procumbens* |
| DG-Sa-A-P2.5 | August | LSM | 2 | 1.279 | tetraploid | *Salicornia procumbens* |
| DG-Sa-A-P3.1 | August | LSM | 3 | 1.281 | tetraploid | *Salicornia procumbens* |
| DG-Sa-A-P3.2 | August | LSM | 3 | 1.261 | tetraploid | *Salicornia procumbens* |
| DG-Sa-A-P3.3 | August | LSM | 3 | 1.322 | tetraploid | *Salicornia procumbens* |
| DG-Sa-A-P4.1 | August | LSM | 4 | 0.633 | diploid | *Salicornia europaea* |
| DG-Sa-A-P4.2 | August | LSM | 4 | 0.695 | diploid | *Salicornia europaea* |
| DG-Sa-A-P4.3 | August | LSM | 4 | 0.634 | diploid | *Salicornia europaea* |
| DG-Sa-A-P5.1 | August | LSM | 5 | 1.342 | tetraploid | *Salicornia procumbens* |
| DG-Sa-A-P5.2 | August | LSM | 5 | 1.278 | tetraploid | *Salicornia procumbens* |
| DG-Sa-A-P5.3 | August | LSM | 5 | 1.270 | tetraploid | *Salicornia procumbens* |
| DG-Sa-A-P5.4 | August | LSM | 5 | 1.299 | tetraploid | *Salicornia procumbens* |
| DG-Sa-A-P6.1 | August | LSM | 6 | 0.622 | diploid | *Salicornia procumbens* |
| DG-Sa-A-P6.2 | August | LSM | 6 | 1.251 | tetraploid | *Salicornia procumbens* |
| DG-Sa-A-P7.1 | August | LSM | 7 | 1.328 | tetraploid | *Salicornia procumbens* |
| DG-Sa-A-P7.2 | August | LSM | 7 | 1.333 | tetraploid | *Salicornia procumbens* |
| DG-Sa-A-P7.3 | August | LSM | 7 | 1.297 | tetraploid | *Salicornia procumbens* |
| DG-Sa-A-P8.1 | August | LSM | 8 | 1.287 | tetraploid | *Salicornia procumbens* |
| DG-Sa-A-P8.2 | August | LSM | 8 | 1.274 | tetraploid | *Salicornia procumbens* |
| DG-Sa-A-P9.1 | August | LSM | 9 | 1.287 | tetraploid | *Salicornia procumbens* |
| DG-Sa-A-P9.2 | August | LSM | 9 | 1.279 | tetraploid | *Salicornia procumbens* |
| DG-Sa-A-P10.1 | August | LSM | 10 | 1.304 | tetraploid | *Salicornia procumbens* |
| DG-Sa-A-P10.2 | August | LSM | 10 | 1.336 | tetraploid | *Salicornia procumbens* |
| DG-Sa-A-P11.1 | August | LSM | 11 | 1.386 | tetraploid | *Salicornia procumbens* |
| DG-Sa-A-P11.2 | August | LSM | 11 | 1.360 | tetraploid | *Salicornia procumbens* |
| DG-Sa-A-P11.3 | August | LSM | 11 | 1.346 | tetraploid | *Salicornia procumbens* |
| DG-Sa-A-P12.1 | August | LSM | 12 | 1.301 | tetraploid | *Salicornia procumbens* |
| DG-Sa-A-P12.3 | August | LSM | 12 | 1.338 | tetraploid | *Salicornia procumbens* |
| DG-Sa-A-P12.4 | August | LSM | 12 | 1.355 | tetraploid | *Salicornia procumbens* |
| DG-Sa-A-P13.1 | August | LSM | 13 | 0.639 | diploid | *Salicornia europaea* |
| DG-Sa-A-P13.2 | August | LSM | 13 | 0.645 | diploid | *Salicornia europaea* |
| DG-Sa-A-P13.3 | August | LSM | 13 | 0.716 | diploid | *Salicornia europaea* |
| DG-Sa-A-P13.4 | August | LSM | 13 | 0.803 | diploid | *Salicornia europaea* |
| DG-Sa-A-P13.5 | August | LSM | 13 | 0.744 | diploid | *Salicornia europaea* |
| DG-Sa-A-P14.1 | August | LSM | 14 | 1.337 | tetraploid | *Salicornia procumbens* |
| DG-Sa-A-P14.2 | August | LSM | 14 | 1.413 | tetraploid | *Salicornia procumbens* |
| DG-Sa-A-P14.3 | August | LSM | 14 | 0.655 | diploid | *Salicornia europaea* |
| DG-Sa-A-P14.4 | August | LSM | 14 | 0.713 | diploid | *Salicornia europaea* |
| DG-Sa-A-P14.5 | August | LSM | 14 | 1.309 | tetraploid | *Salicornia procumbens* |
| DG-Sa-A-P15.1 | August | LSM | 15 | 0.609 | diploid | *Salicornia europaea* |
| DG-Sa-A-P15.2 | August | LSM | 15 | 1.388 | tetraploid | *Salicornia procumbens* |
| DG-Sa-A-P15.3 | August | LSM | 15 | 0.694 | diploid | *Salicornia europaea* |
| DG-Sa-A-P15.4 | August | LSM | 15 | 0.683 | diploid | *Salicornia europaea* |
| DG-Sa-A-P16.1 | August | LSM | 16 | 1.263 | tetraploid | *Salicornia procumbens* |
| DG-Sa-A-P16.2 | August | LSM | 16 | 1.422 | tetraploid | *Salicornia procumbens* |
| DG-Sa-A-P16.3 | August | LSM | 16 | 1.265 | tetraploid | *Salicornia procumbens* |
| DG-Sa-A-P17.1 | August | LSM | 17 | 0.599 | diploid | *Salicornia europaea* |
| DG-Sa-A-P17.2 | August | LSM | 17 | 0.662 | diploid | *Salicornia europaea* |
| DG-Sa-A-P18.1 | August | LSM | 18 | 0.601 | diploid | *Salicornia europaea* |
| DG-Sa-A-P18.2 | August | LSM | 18 | 0.627 | diploid | *Salicornia europaea* |
| DG-Sa-A-P19.1 | August | LSM | 19 | 1.337 | tetraploid | *Salicornia procumbens* |
| DG-Sa-A-P19.2 | August | LSM | 19 | 0.648 | diploid | *Salicornia europaea* |
| DG-Sa-A-P20.1 | August | LSM | 20 | 1.292 | tetraploid | *Salicornia procumbens* |
| DG-Sa-A-P20.2 | August | LSM | 20 | 1.323 | tetraploid | *Salicornia procumbens* |
| DG-Sa-A-P21.1 | August | LSM | 21 | 1.302 | tetraploid | *Salicornia procumbens* |
| DG-Sa-A-P21.2 | August | LSM | 21 | 1.281 | tetraploid | *Salicornia procumbens* |
| DG-Sa-A-P21.3 | September | LSM | 21 | 0.605 | diploid | *Salicornia europaea* |

**Table S3**. Distribution of *S. europaea* and *S. procumbens* collected for root anatomical studies.

| **Sample ID** | **Sampling point** | **Genome Size (2C DNA content, pg)** | **Ploidy** | **Species** |
| --- | --- | --- | --- | --- |
| LSM1 | 1 | 0.705 | diploid | *Salicornia europaea* |
| LSM2 | 1 | 1.385 | tetraploid | *Salicornia procumbens* |
| LSM3 | 2 | 1.383 | tetraploid | *Salicornia procumbens* |
| LSM4 | 2 | 1.278 | tetraploid | *Salicornia procumbens* |
| LSM5 | 3 | 1.349 | tetraploid | *Salicornia procumbens* |
| LSM6 | 3 | 0.659 | diploid | *Salicornia europaea* |
| LSM7 | 4 | 0.644 | diploid | *Salicornia europaea* |
| LSM8 | 4 | 0.686 | diploid | *Salicornia europaea* |
| LSM9 | 5 | 1.285 | tetraploid | *Salicornia procumbens* |
| LSM10 | 5 | 0.684 | diploid | *Salicornia europaea* |
| LSM11 | 6 | 0.718 | diploid | *Salicornia europaea* |
| LSM18 | 6 | 1.384 | tetraploid | *Salicornia procumbens* |
| LSM19 | 7 | 1.292 | tetraploid | *Salicornia procumbens* |
| LSM21 | 7 | 1.259 | tetraploid | *Salicornia procumbens* |
| LSM22 | 8 | 1.211 | tetraploid | *Salicornia procumbens* |
| LSM23 | 8 | 1.292 | tetraploid | *Salicornia procumbens* |
| LSM24 | 9 | 0.608 | diploid | *Salicornia europaea* |
| LSM25 | 9 | 1.239 | tetraploid | *Salicornia procumbens* |
| LSM26 | 10 | 1.231 | tetraploid | *Salicornia procumbens* |
| LSM27 | 10 | 1.245 | tetraploid | *Salicornia procumbens* |
| LSM28 | 11 | 1.345 | tetraploid | *Salicornia procumbens* |
| LSM29 | 11 | 0.641 | diploid | *Salicornia europaea* |
| LSM30 | 12 | 0.603 | diploid | *Salicornia europaea* |
| LSM31 | 12 | 0.592 | diploid | *Salicornia europaea* |
| LSM32 | 13 | 1.292 | tetraploid | *Salicornia procumbens* |
| LSM33 | 13 | 1.295 | tetraploid | *Salicornia procumbens* |
| LSM34 | 14 | 1.344 | tetraploid | *Salicornia procumbens* |
| LSM35 | 14 | 1.249 | tetraploid | *Salicornia procumbens* |
| LSM36 | 15 | 1.313 | tetraploid | *Salicornia procumbens* |
| LSM37 | 15 | 1.361 | tetraploid | *Salicornia procumbens* |
| LSM38 | 16 | 0.654 | diploid | *Salicornia europaea* |
| LSM39 | 16 | 1.264 | tetraploid | *Salicornia procumbens* |
| LSM43 | 17 | 1.313 | tetraploid | *Salicornia procumbens* |

**Table S4.** Different indices of alpha diversity estimated to compare samples of each cytotype collected in spring and summer. (Kruskal-Wallis test, p < 0.05).

| ***Alpha Diversity Index*** | ***Sample*** | ***Sample*** | ***H*** | ***p-value*** | ***q-value*** |
| --- | --- | --- | --- | --- | --- |
| Shannon Diversity | diploid-spring(n=8) | diploid-summer(n=10) | 0.386842 | 0.533964 | 0.533964 |
| Shannon Diversity | tetraploid-spring(n=5) | tetraploid-summer(n=7) | 2.380220 | 0.122880 | 0.147456 |
| Pielou's evenness | diploid-spring (n=8) | diploid-summer(n=10) | 0.639474 | 0.423901 | 0.423901 |
| Pielou's evenness | tetraploid-spring(n=5) | tetraploid-summer(n=7) | 2.380220 | 0.122880 | 0.147456 |
| Observed richness | diploid-spring (n=8) | diploid-summer(n=10) | 0.579971 | 0.446324 | 0.535588 |
| Observed richness | tetraploid-spring(n=5) | tetraploid-summer(n=7) | 0.110115 | 0.740014 | 0.740014 |
| Faith's Phylogenetic Diversity | diploid-spring (n=8) | diploid-summer(n=10) | 0.000000 | 1.000000 | 1.000000 |
| Faith's Phylogenetic Diversity | tetraploid-spring(n=5) | tetraploid-summer(n=7) | 0.006593 | 0.935283 | 1.000000 |

**Table S5.** Functional assignment for each fungal genera detected in roots of *S. europaea* and *S. procumbens*. Na: information not available.

|  |  |  | **FunGuild database** | **FungalTrait database** | | | |
| --- | --- | --- | --- | --- | --- | --- | --- |
| **Host** | **Phylum** | **Genus** | **Trophic mode** | **Primary lifestyle** | **Secondary lifestyle** | **Endophytic capability** | **Comments of lifestyle** |
| *S. procumbens* | Ascomycota | *Lasiodiplodia* | Pathotroph | plant pathogen | foliar endophyte | foliar endophyte | na |
| *S. procumbens* | Ascomycota | *Didymella* | Pathotroph-Saprotroph | plant pathogen | litter saprotroph | na | hypervariable |
| *S. procumbens* | Ascomycota | *Ophiosphaerella* | Saprotroph | plant pathogen | litter saprotroph | no endophytic capacity | na |
| *S. procumbens* | Basidiomycota | *Cystofilobasidium* | Saprotroph | litter saprotroph | na | na | na |
| *S. procumbens* | Basidiomycota | *Solicoccozyma* | na | soil saprotroph | epiphyte | na | na |
| *S. procumbens* | Basidiomycota | *Tausonia* | na | soil saprotroph | na | na | na |
| *S. europaea* | Ascomycota | *Neodidymelliopsis* | na | litter saprotroph | na | na | na |
| *S. europaea* | Ascomycota | *Preussia* | Saprotroph | dung saprotroph | na | na | na |
| *S. europaea* | Ascomycota | *Trematosphaeria* | Saprotroph | wood saprotroph | na | no endophytic capacity | na |
| *S. europaea* | Ascomycota | *Acremonium* | Pathotroph-Saprotroph-Symbiotroph | unspecified saprotroph | foliar endophyte | foliar endophyte | hypervariable |
| *S. europaea* | Basidiomycota | *Kurtzmanomyces* | na | unspecified saprotroph | na | na | na |
| *S. europaea* | Basidiomycota | *Microstroma* | Pathotroph | plant pathogen | na | no endophytic capacity | na |
| *S. europaea* | Basidiomycota | *Rhodotorula* | Pathotroph-Saprotroph | unspecified saprotroph | foliar endophyte | foliar endophyte | na |
| *S. europaea* | Basidiomycota | *Sporobolomyces* | Pathotroph-Saprotroph | mycoparasite | na | na | some species litter saprotroph |
| *S. europaea* | Basidiomycota | *Itersonilia* | na | plant pathogen | na | na | na |
| *S. europaea* | Basidiomycota | *Papiliotrema* | na | mycoparasite | fungal decomposer | na | na |
| both | Ascomycota | *Cladosporium* | na | litter saprotroph | plant pathogen | foliar endophyte | na |
| both | Ascomycota | *Mycosphaerella* | Pathotroph | plant pathogen | litter saprotroph | na | na |
| both | Ascomycota | *Laburnicola* | na | wood saprotroph | na | no endophytic capacity | na |
| both | Ascomycota | *Paraphaeosphaeria* | Saprotroph | wood saprotroph | na | no endophytic capacity | na |
| both | Ascomycota | *Lentithecium* | na | wood saprotroph | na | no endophytic capacity | na |
| both | Ascomycota | *Byssothecium* | Saprotroph | unspecified saprotroph | na | na | na |
| both | Ascomycota | *Alternaria* | Pathotroph-Saprotroph-Symbiotroph | plant pathogen | litter saprotroph | foliar endophyte | na |
| both | Ascomycota | *Neocamarosporium* | na | wood saprotroph | plant pathogen | na | no endophytic capacity |
| both | Ascomycota | *Aspergillus* | na | unspecified saprotroph | foliar endophyte | foliar endophyte | hypervariable, saprotroph, animal pathogen, endophyte |
| both | Ascomycota | *Plectosphaerella* | na | plant pathogen | litter saprotroph | na | na |
| both | Ascomycota | *Sarocladium* | Saprotroph | plant pathogen | litter saprotroph | na | na |
| both | Ascomycota | *Fusarium* | Pathotroph-Saprotroph-Symbiotroph | plant pathogen | litter saprotroph | foliar endophyte | hypervariable, saprotroph, damping-off pathogen |
| both | Ascomycota | *Gibberella* | Pathotroph | plant pathogen | litter saprotroph | foliar endophyte | hypervariable, saprotroph , damping-off pathogen |
| both | Ascomycota | *Ilyonectria* | Pathotroph | plant pathogen | na | na | na |
| both | Ascomycota | *Slopeiomyces* | na | root endophyte | na | root endophyte | na |
| both | Ascomycota | *Microdochium* | na | plant pathogen | foliar endophyte | foliar endophyte | na |
| both | Basidiomycota | *Botryobasidium* | na | wood saprotroph | na | na | na |
| both | Basidiomycota | *Bensingtonia* | Saprotroph | unspecified saprotroph | na | na | na |
| both | Basidiomycota | *Malassezia* | na | soil saprotroph | root-associated | root endophyte | na |
| both | Basidiomycota | *Filobasidium* | Saprotroph | unspecified saprotroph | na | na | na |
| both | Basidiomycota | *Naganishia* | na | unspecified saprotroph | na | na | extremophile |
| both | Basidiomycota | *Dioszegia* | na | litter saprotroph | na | na | na |
| both | Basidiomycota | *Vishniacozyma* | na | soil saprotroph | na | na | extremophile |
| both | Basidiomycota | *Wallemia* | Saprotroph | unspecified saprotroph | na | na | extremophile for high salt or sugar content, xerophile |
